# Secure and federated linear mixed model association tests

**DOI:** 10.1101/2022.05.20.492837

**Authors:** Jeffrey Chen, Manaswitha Edupalli, Bonnie Berger, Hyunghoon Cho

## Abstract

Privacy-preserving algorithms for genome-wide association studies (GWAS) promise to facilitate data sharing across silos to accelerate new discoveries. However, existing approaches do not support an important, prevalent class of methods known as linear mixed model (LMM) association tests or would provide limited privacy protection, due to the high computational burden of LMMs under existing secure computation frameworks. Here we introduce SafeGENIE, an efficient and provably secure algorithm for LMM-based association studies, which allows multiple entities to securely share their data to jointly compute association statistics without leaking any intermediary results. We overcome the computational burden of LMMs by leveraging recent advances in LMMs and secure computation, as well as a novel scalable dimensionality reduction technique. Our results show that SafeGENIE obtains accurate association test results comparable to a state-of-the-art centralized algorithm (REGENIE), and achieves practical runtimes even for large datasets of up to 100K individuals. Our work unlocks the promise of secure and distributed algorithms for collaborative genomic studies.^1^

## 1 Introduction

Genome-wide association studies (GWAS) have been a major driving force behind genomics research [1,2]. Ongoing efforts to amass large and diverse collections of genomic data, such as the All of Us Program [3] and the UK Biobank [4], will continue to bring new discoveries of genetic variants of biological and clinical importance. However, many of the existing genomic datasets are held in isolated repositories within national or organizational boundaries under strict data sharing limitations, presenting a key hurdle for collaborative research [4-6]. Achieving sufficient statistical power for rare diseases and underrepresented populations (e.g. admixed individuals) requires new strategies to facilitate sharing of genomic data across multiple data repositories.

To this end, recent studies have proposed a range of privacy-preserving solutions for GWAS. These works aim to allow analyses to be jointly performed on multiple parties’ datasets without sharing the raw individual-level data, thus providing a path to circumvent regulatory restrictions. In particular, cryptographic approaches based on secure computation frameworks [7–10], such as homomorphic encryption (HE) [11, 12] and secure multiparty computation (MPC) [13], offer the strongest notions of data privacy, which state that any (encrypted) data that is externally shared by each party is statistically indistinguishable from random (under certain security assumptions, such as the parties do not collude in the case of MPC) [14]. Alternative approaches based on trusted execution environment (TEE) technology [15, 16] or distributed/federated algorithms [17, 18] have also been proposed, albeit providing weaker forms of privacy protection.

A key limitation shared by most existing solutions for privacy-preserving GWAS is that they consider simplified analysis workflows that do not reflect the current standard practice in genomics. Specifically, population stratification correction—a crucial step in GWAS for mitigating confounding effects arising from the implicit association between population structure in the study cohort and the target phenotype [19]— has long been unaddressed by solutions for privacy-preserving GWAS; the one exception is recent work that incorporated an approach based on principal components analysis (PCA) [13]. The gap between current solutions and practical workflows is in part due to the fact that existing techniques for privacy protection typically incur a computational overhead that grows with the complexity of the algorithm being implemented; this renders the task of making these sophisticated algorithms secure for modern GWAS highly challenging. Overcoming this barrier is a necessary step to realize the full potential of emerging privacy techniques.

*Linear mixed models* (LMMs) are one of two main classes of methods in the literature for addressing population stratification in GWAS. The other is a more traditional PCA-based approach [20], where the top principal components of the genotype matrix are included as fixed-effect covariates in the statistical models of genetic association to control for global population structure. In contrast, LMMs view the ancestry-related effect on the phenotype as a *random* effect, whose covariance is determined by the genetic relatedness patterns in the study cohort, also estimated from the data. LMMs are known to be more effective at capturing cryptic relatedness and fine-grain population structure within the study cohort [21]. Further enabled by recent algorithmic advances on reducing the computational burden of LMM computation [21–23], LMMs are increasingly becoming the preferred approach for GWAS [24]. However, little work exists in the literature on *privacy-preserving* LMM-based GWAS over decentralized, or distributed, datasets, which we ascribe to the complexity of existing LMM algorithms and their high computational cost even with full access to the data. This severe bottleneck hinders the use of privacy-preserving techniques for a full GWAS pipeline in practice.

In this work, we present SafeGENIE (secure and federated REGENIE), a privacy-preserving algorithm for performing LMM-based association tests on distributed genomic datasets. We develop a scalable algorithm for privacy-preserving LMM computation by synthesizing recent advances in cryptography, distributed algorithms, and population genetics, including: multiparty homomorphic encryption [25], distributed linear regression models [26], an efficient stacked ridge regression approach for LMMs (called REGENIE [27]). SafeGENIE provides formal security guarantees offered by the underlying cryptographic frameworks, including protection for all intermediate results, while maintaining efficiency by maximally leveraging local computation using plaintext (raw) data. Our work also incorporates new algorithmic strategies (e.g. linear algebra techniques for low-rank matrix update) to overcome the unique computational bottlenecks that arise when jointly utilizing the aforementioned techniques in SafeGENIE. To our knowledge, SafeGENIE is the first privacy-preserving algorithm to fully implement the LMM-based GWAS pipeline.

Our experimental results show that SafeGENIE produces LMM association statistics closely matching those of the plaintext algorithm REGENIE, demonstrating its utility for real-world studies. SafeGENIE also maintains runtimes on the order of days for datasets including up to a hundred thousand individuals and is efficient in memory usage. Although this runtime reflects the heavy computational burden of the underlying LMM computation, we demonstrate that our method is a significant improvement over the state-of-the-art baseline approaches and can be readily applied to many existing datasets. Our work represents an important step towards enabling a wide range of genomic analyses to be performed while protecting the private data.

## 2 Problem Definitions and Review of Existing Methods

### 2.1 Linear Mixed Model (LMM) Association Studies

We start by formally describing the LMMs used for genome-wide association tests. LMMs model the target phenotype vector **y** of length *N* individuals using the following linear model

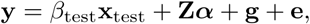

where **x**_test_ is a vector of allele dosages of the variant being tested across *N* individuals, **Z** is a *N*-by-*C* matrix of observed covariates where *C* denotes the number of covariates, **g** represents the ambient genetic effect, and **e** represents the environmental effect. Both **x**_test_ and **y** are standardized to have zero mean and unit variance. We let *N* and *M* be the number of individuals and the number of variants in the dataset, respectively.

In this model, *β*_test_ and *α* describe the fixed effect sizes associated with the tested variant and the covariates, respectively, whereas **g** and **e** are modeled as random effects (hence the term “mixed” model). Under the standard infinitesimal model, which posits that the genetic effect on phenotype consists of many small effect-size variants, we can express

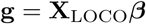

where **X**_LOCO_ is a *N*-by-*M*_LOCO_ matrix consisting of the standardized genotypes of *M*_LOCO_ variants used to model the genetic effect based on the standard leave-one-chromosome-out (LOCO) scheme, which excludes all variants in the same chromosome as the tested variant in order to avoid the effects of linkage disequilibrium. These ambient variants are associated with effect sizes 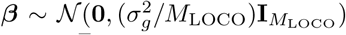, inducing a distribution over the genetic effect as 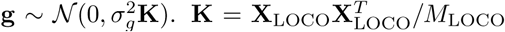 is referred to as the genetic relatedness (or empirical kinship) matrix. The environmental effect is modeled as 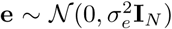. Note that *σ*_*g*_ and *σ*_*e*_ represent the variances of the polygenic and environmental components. The goal of the association test is to test the null hypothesis *H*_0_ : *β*_test_ = 0.

A standard technique [21, 27] is to project the covariates out of the phenotypes and genotypes to simplify the computation. This results in a modified model

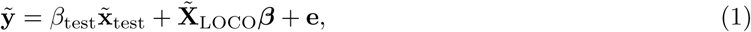

where 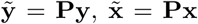, and 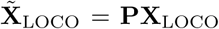 with **P** = **I**_*N*_ – **Z**(**Z**^*T*^**Z**)^*-*1^**Z**^*T*^. Note that **P** projects a vector onto the null space of **Z**.

The LMM-based χ^2^ test statistic is given by

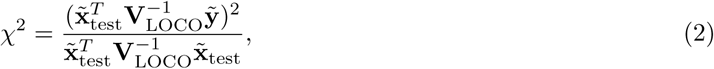

where 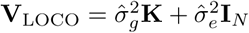 given the estimates 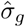 and 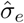 of the variance parameters.

### 2.2 Review of REGENIE: Efficient Stacked Regression for LMMs

Computing the LMM association statistics is a computationally expensive task, in part because the maximum likelihood estimation of the variance parameter *σ*_*g*_ involves costly matrix operations involving the N-by-N genetic relatedness matrix (GRM), which becomes prohibitively large for large-scale datasets. Much of the prior algorithmic development efforts have focused on speeding up the use of GRM, e.g. by exploiting a factorization of the matrix [23]. A recent algorithm called REGENIE [27] introduced a different strategy based on *stacked regression*, resulting in significant scalability improvements for LMM-based association tests, while achieving comparable accuracy to existing state-of-the-art tools such as BOLT-LMM [21], fastGWA [22], and SAIGE [28]. Indeed, we observed similar performance between REGENIE and BOLT-LMM on the benchmark dataset used in our study (Supplementary Figure 1). Moreover, we discovered that REGENIE’s approach is more amenable to efficient computation over distributed datasets, thereby forming an important basis for our SafeGENIE method.

In REGENIE [27], the whole-genome regression model in Equation 1 is estimated in two phases by first regressing out the contribution of 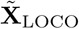 from 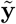, then fitting *β*_test_ on the residuals to test for association. To further reduce the cost of regression over the large genome-wide matrix 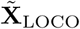, REGENIE employs a stacked regression approach in the following two steps, referred to as Levels 0 and 1.

**Level 0**. The genotype matrix is first split into *B* contiguous blocks of *T* variants:

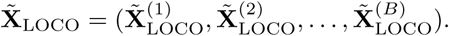

Then for each block *b* ∈ [*B*] and different choices of the regularization parameter *λ*_*r*_ *∈* {*λ*_1_, …, *λ*_*R*_}, where *R* represents the number of regularization parameters being tested, the following solution to the ridge regression problem 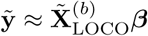 is computed.

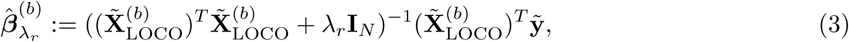

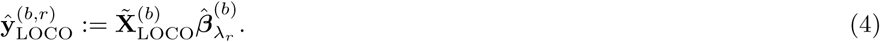

The 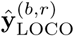 is referred to as the *predictors*, representing the best polygenic prediction of the phenotype within a given genomic region, accounting for genotype correlations.

**Level 1**. The local predictors from Level 0 are aggregated to form a *N*-by-*BR* global feature matrix

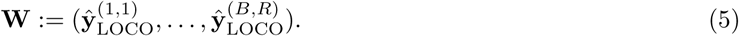

Then another around of ridge regression is performed (with K-fold cross validation to choose the optimal regularization parameter *η*) to obtain the following genome-wide phenotype predictions:

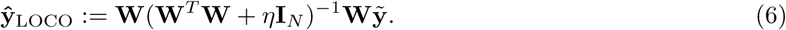

Given this global predictor as a proxy for ambient genetic effect, the *χ*^2^ statistic (with one degree of freedom) for the variant being tested in Equation 2 can now be formulated as

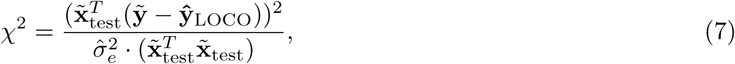

where 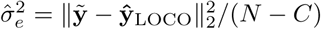is the estimated variance of the environmental effect.

The above approach substantially improves the speed of LMM computation by decomposing the problem into *B* separate local ridge regression problems, which can be performed in parallel in high performance computing environments. Importantly, by formulating the problem as a series of ridge regression tasks, the problem becomes more tractable for secure distributed computation, an aspect we exploit in our design of SafeGENIE to achieve efficiency.

### 2.3 Review of Privacy-Preserving GWAS Algorithms and Their Limitations

Prior works have developed privacy-aware algorithms for conducting GWAS over private datasets to facilitate genomic data sharing and collaboration. These methods leverage a range of different computational techniques with different strengths and weaknesses. Methods based on cryptographic frameworks such as secure multiparty computation (MPC) [7, 29] and homomorphic encryption (HE) [9, 30] aim to directly allow computation over encrypted data. MPC relies on interactive protocols among multiple parties to carry out the joint computation without revealing the private data, whereas HE enables non-interactive computation over the ciphertexts, albeit incurring higher computational overhead. GWAS algorithms based on both approaches have been developed [11–13], but they do not support LMM-based analyses.

Another branch of methods follow a distributed or federated algorithm design [17, 18], leveraging local computation performed by each party on their respective dataset combined with aggregation steps that exchange information among the parties to move towards a global solution. Although these methods enjoy higher computational flexibility and efficiency compared to cryptographic approaches, there is a general lack of understanding of the extent to which private information is leaked in the intermediate results shared among the parties, thus providing limited privacy protection. Notably, a recent work in this line presented a promising solution for distributed training of generalized LMMs [31], which is closely related to our work. However, we note that the prior method did not address association testing and supports only a single covariate, thus the LMM-based GWAS remains an open problem. Approaches based on trusted execution environments, such as the Intel SGX enclave, have also been proposed [15,16]. They perform computation on private data securely within a secure hardware component that is isolated from the rest of the host operating system. Similar to the plaintext distributed approach, these methods are efficient but offer limited privacy protection due to various security vulnerabilities that security researchers continue to discover, necessitating careful mitigation strategies (e.g. [32]). In this work, we adopt the cryptographic paradigm for secure computation, which offers the strongest notion of privacy, and focus on efficient algorithm design to address the associated computational burden.

## 3 Our Method: SafeGENIE

### 3.1 Overview of SafeGENIE

SafeGENIE is a privacy-preserving algorithm for LMM association analysis, which jointly analyzes private datasets held by different entities without leaking individual-level information. The results of SafeGENIE, by design, closely match those of running REGENIE on a pooled dataset with minimal loss in precision, while additionally protecting data privacy. The core computational framework of SafeGENIE is a hybrid of secure multiparty computation (MPC) and homomorphic encryption (HE), building upon the recently introduced framework of multiparty homomorphic encryption (MHE) [25,33–35]. Combining the ability of HE to perform non-interactive secure computation at each site with efficient MPC protocols for high-complexity operations such as comparison and inverse square root [13], our hybrid approach enables SafeGENIE to leverage efficient local plaintext computation while maintaining flexibility and accuracy in computational capabilities. In addition, our framework provides formal privacy guarantees by only exchanging *encrypted* forms of all intermediate results in accordance with established HE and MPC techniques throughout the protocol to facilitate the distributed computation (in contrast to other federated approaches that leak intermediate results and therefore sensitive data). Using these cryptographic tools, SafeGENIE implements our distributed algorithm for stacked regression for LMMs, inspired by the success of the centralized plaintext algorithm introduced by REGENIE. We introduce new algorithmic techniques for maximizing local computation that allow SafeGENIE to achieve near-constant scalability to larger datasets as we show in our results. We provide a graphical illustration of SafeGENIE in Figure 1.

**Figure 1:**
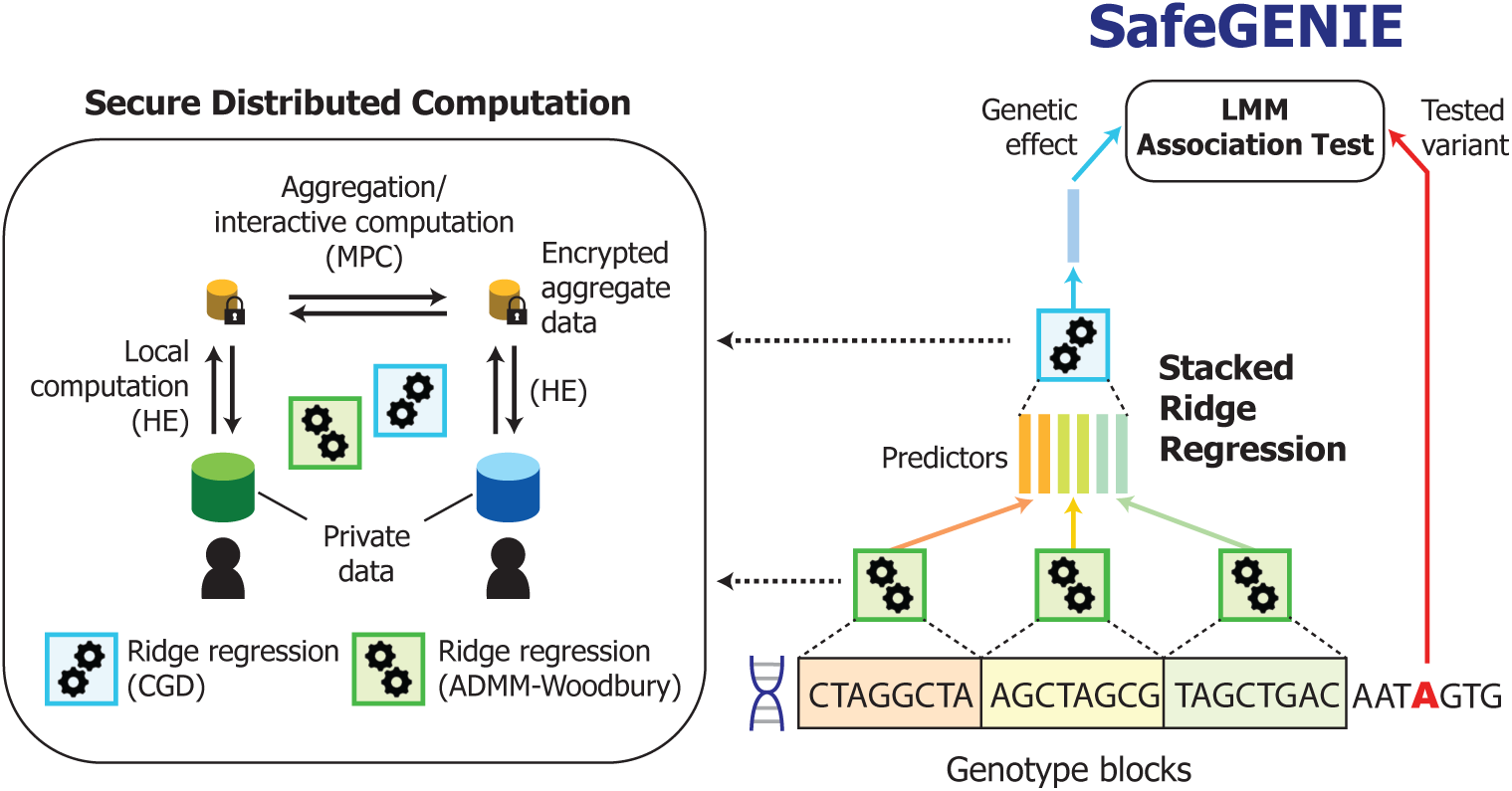
Overview of SafeGENIE. SafeGENIE implements a stacked ridge regression approach [27] to LMM association testing, whereby regression models trained on local genomic blocks are combined in another round of regression to produce estimated background genetic effects for the association tests. **(Left)** Both levels of ridge regression are performed using secure distributed algorithms we developed (CGD and ADMM-Woodbury), which leverage both homomorphic encryption and secure multiparty computation frameworks to provide privacy protection.

### 3.2 Our Secure Computation Model

In our application setting, we consider *P* independent parties, each with a local GWAS dataset, who wish to jointly perform LMM association analysis on the pooled data without revealing private data. Our core approach is to keep these local datasets in plaintext and to encrypt, using MHE, only the intermediate results that need to be aggregated across the parties. MHE [25] is an extension of homomorphic encryption (HE) schemes where the decryption key is split among multiple parties using secret sharing, which ensures that encryption and homomorphic operations (performing computation over the secrets without decrypting the data) can be performed locally, while decryption requires all parties to cooperate, thus giving each party control over which pieces of results (such as the final association statistics) are revealed to other parties.

Importantly, MHE enables our distributed approach to secure computation, where we iterate between (1) a local computation phase, where each party computes local results leveraging both local plaintext data and encrypted shared data, and (2) an aggregation phase, where the parties interact to update the encrypted shared data, to perform the desired analysis. We additionally incorporate secret sharing-based MPC routines into our protocol by securely switching data representation between HE and secret shares. Secret shared data allow us to leverage efficient interactive protocols for more complex operations at the expense of higher communication. In our protocols, we securely perform comparison, inverse square root, and eigendecomposition over a small matrix using interactive MPC subroutines, while relying on HE for the rest. Note that, following the prior work on MPC-based GWAS [13], we adopt an efficient server-aided model for MPC, in which a coordinating party facilitates the computation by supplying the main parties with random numbers satisfying a certain structure. The coordinating party does not receive any portion of the private data other than the knowledge of data dimensions.

For the purposes of describing our distributed LMM algorithm implemented in SafeGENIE, we view our secure computation framework as a collection of building block protocols, each implementing a simple operation such as matrix multiplication or square root, which we compose to implement an end-to-end secure protocol that implements the LMM computation. As the focus of this work is on distributed algorithm design, in particular on optimizing the use of secure subroutines to efficiently perform LMM computation, we refer to relevant prior works on MPC and MHE for detailed descriptions of the subroutines used in SafeGENIE [12, 13].

### 3.3 Key Challenges of Distributed LMM Computation

To motivate our novel algorithmic techniques, we first describe the computational challenges in distributing the LMM computation. Recall that state-of-the-art LMM-based GWAS algorithms (e.g. BOLT-LMM [21]) account for population stratification by using the N-by-N genetic relatedness matrix (GRM) **K**, which intuitively captures how individuals within a dataset are related to one another. A core computational step in the estimation of variance parameters or the calculation of association statistics is calculating a quantity of the form

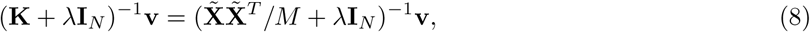

for some length-N vector **v**. Because the size of **K** scales with the number of individuals in the dataset, for large-scale GWAS, this computation inherently incurs an overwhelming computational cost. As such, addressing this challenge has been the focus of recent algorithmic development efforts for LMM.

In our setting, the fact that the off-diagonal blocks of **K** describe relatedness between individuals in *different* collaborating sites introduces a unique difficulty in distributing the computation. When naïvely implemented, those interactive terms in **K** are bound to require heavy communication among parties to account for their contributions. Even recently proposed iterative approaches for efficiently solving this linear system of equations without the inverse (e.g. conjugate gradient descent used by BOLT-LMM [21]) presents a similar challenge, as it involves repeated multiplications of **K** with a candidate solution vector. Moreover, we note that **K** is typically defined over covariate-corrected genotypes 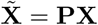, where **P** denotes a projection matrix for removing the covariate effect, which introduces another layer of entanglement between the private datasets at different sites, making distributed computation further challenging.

### 3.4 Our Approach: Secure Distributed Ridge Regression with Covariates

SafeGENIE approaches this problem differently. Following the methodology of REGENIE [27], instead of using the notion of a GRM, we train ridge regression models for local genomic windows to use as a proxy for ambient genetic effect on phenotype. Correcting for these polygenic predictions by testing for a variant’s association with the phenotype residuals, our approach implicitly accounts for population stratification under the LMM formulation.

Our work identifies this alternative approach for LMMs as a key enabling factor for distributing the computation. Let 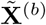 be a subset of columns from a genotype matrix corresponding to a genomic block with *M*_*b*_ variants. The ridge regression problem for LMM reduces to computing the expression

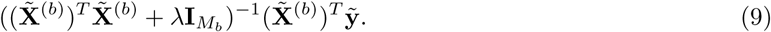

In contrast to Equation 8, we see that the inverse operation is for a matrix of size *M*_*b*_-by-*M*_*b*_, which is significantly smaller than **K** (note *M*_*b*_ is typically 1000). Furthermore, for horizontally distributed 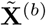 where each party p holds a subset of rows in this matrix denoted 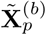, we have the following decomposition

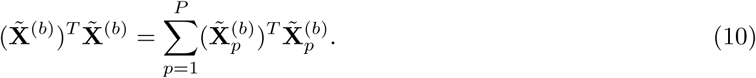

This property allows SafeGENIE to distribute the computation more efficiently across the parties and maximally leverage plaintext data that is available locally. In the following section, we describe how we exploit this insight to design secure and distributed algorithms for conjugate gradient descent (CGD) and alternating direction method of multipliers (ADMM) approaches for ridge regression, which are used by SafeGENIE to carry out the Level 1 and Level 0 steps of REGENIE, respectively.

### 3.5 SafeGENIE Algorithm Details

Recall from Section 2.2 that stacked regression approach to LMM involves solving the following two ridge regression problems in Levels 0 and 1.

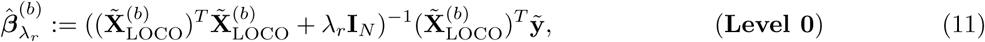

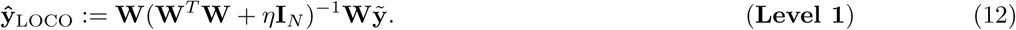

The first is solved *KBR* times for each pair of a block *b* ∈ [*B*] and a regularization parameter, *λ*_*r*_ for *r* ∈ [*R*] using *K*-fold cross validation, and the second is solved *KR* times for each of *R* values of η using the same *K*-fold cross validation. The challenging step in both is the multiplication by the inverse matrix, which is infeasible to solve directly when the matrix is only available in encrypted form. This is unlike REGENIE, which explicitly solves for the inverse using eigenfactorization. Explicitly computing the inverse for a large, homomorphically encrypted matrix imposes a considerable computational burden, which we aim to avoid. To address this, we develop two algorithms described below.

#### 3.5.1 Secure Distributed Conjugate Gradient Descent (CGD)

Conjugate Gradient Descent (CGD) [36] is a well-known iterative algorithm for solving a system of linear equations without explicitly constructing the inverse of the design matrix. Notably, BOLT-LMM heavily utilizes the CGD algorithm to avoid working with the inverse of GRM. A requirement of CGD is that the design matrix be positive definite and well conditioned; in our setting, the regularization term in ridge regression ensures this property [37]. Hence, CGD can be applied to any of the ridge regression problems in our task. The central step in CGD is a multiplication of a candidate solution vector with the design matrix (not the inverse), which lends itself to efficient distributed computation. We outline our distributed CGD algorithm in Algorithm 1.

##### Algorithm 1

Distributed Conjugate Gradient Descent for Ridge Regression

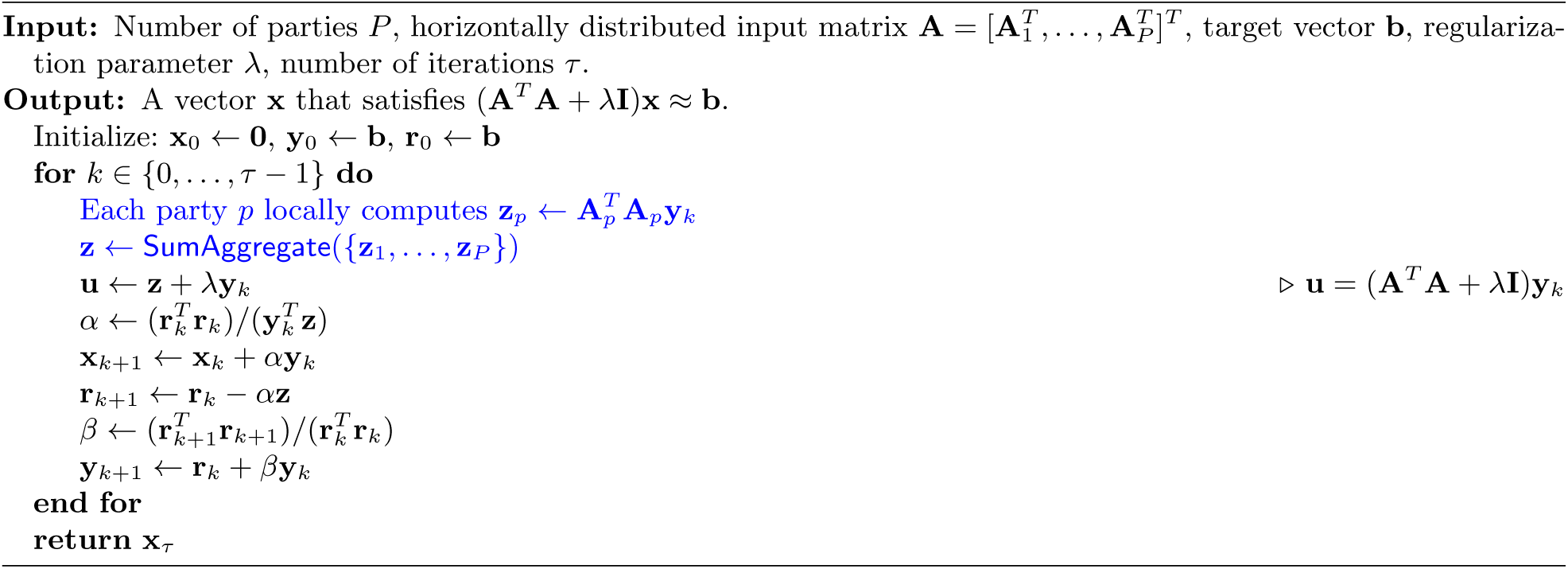

We highlighted in blue the modified step in the distributed approach. As explained in Section 3.4, the fact that the design matrix **A**^*T*^ **A** in our setting decomposes as a sum of local design matrices 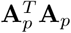 allows this step to be performed independently, then aggregated after both multiplications (the SumAggregate step). Thus, the required communication scales with the number of predictive features (variants), not the number of samples held by each party, thereby offering better scaling to large datasets. To apply this algorithm to encrypted datasets, each of the calculations, namely matrix-vector multiplication, inner products, and addition/subtraction, are implemented using HE routines, with the exception of division, for which we switch to a secret sharing-based MPC routine. Note that the SumAggregate step involves each party broadcasting their share to others and adding up all shares (homomorphically), which does not involve any decryption. For efficiency, this procedure is implemented over a star network where a central coordinator aggregates all data and in turn relays the result to all parties.

#### 3.5.2 Secure ADMM for Ridge Regression with Covariates

Although CGD offers a natural distributed solution for ridge regression, it is still computationally burdensome for large input matrices. For instance, consider applying CGD to Level 0 of REGENIE, where the input is given as

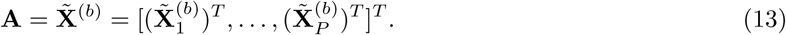

When each party performs the following local computation in Algorithm 1

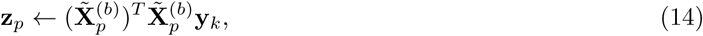

they first need to multiply **y**_*k*_ with 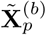, which has an output dimension of *N*_*p*_ (number of individuals in party p’s dataset), followed by another multiplication with 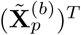, finally resulting in a vector of length *M* _*b*_. This is due to the fact that 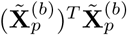 cannot be precomputed in plaintext, since 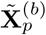 requires covariate correction involving all parties covariate data. Note that *M*_*b*_, the block size, is a user parameter typically set to a small value (e.g. 1000) whereas *N*_*p*_ can grow much larger for large-scale datasets. Therefore, CGD does not benefit from any dimension reduction (to *M*_*b*_) that is otherwise exhibited in the plaintext formulation.

SafeGENIE overcomes this challenge by leveraging the alternating direction method of multipliers (ADMM) technique [38], which is a powerful method for transforming convex optimization problems into distributed optimization problems that can be more efficiently solved. Intuitively, ADMM relaxes the global objective by decoupling the terms involving each individual dataset, which in turn can be jointly optimized using local update equations that, in our case, involve plaintext matrices of size *M*_*b*_ as desired. Our techniques draw inspiration from a recent work in security literature, which introduced a secure multiparty ADMM algorithm for distributed linear regression [26]. Our work extends this work to the setting where the design matrix must be covariate-corrected, which introduces additional challenges as we describe below.

Here we describe how we apply ADMM to the ridge regression problem in Level 0 of REGENIE. Recall that the ridge regression of 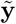 onto 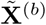 with regularization parameter λ can be equivalently formulated as the following optimization problem:

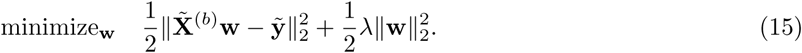

To apply ADMM, we first decouple the two terms using a slack variable **z** with an equality constraint as follows.

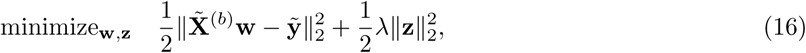

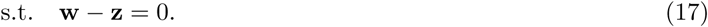

Next, we note that the first objective term can be written as a sum of squared loss computed over each party’s dataset as

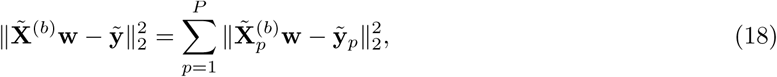

where we partition 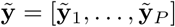 in the same manner as 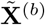. Finally, further decoupling the **w** parameters across parties for distributed optimization, we obtain

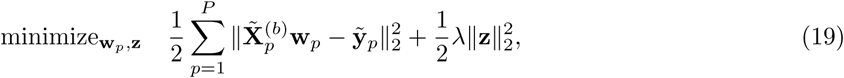

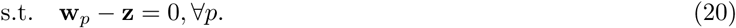

The resulting iterative optimization procedure based on the standard ADMM derivation, for general local matrices **A**_1_,…, **A**_*P*_, is shown in Algorithm 2.

##### Algorithm 2

Standard ADMM Algorithm for Ridge Regression (adapted from [38])

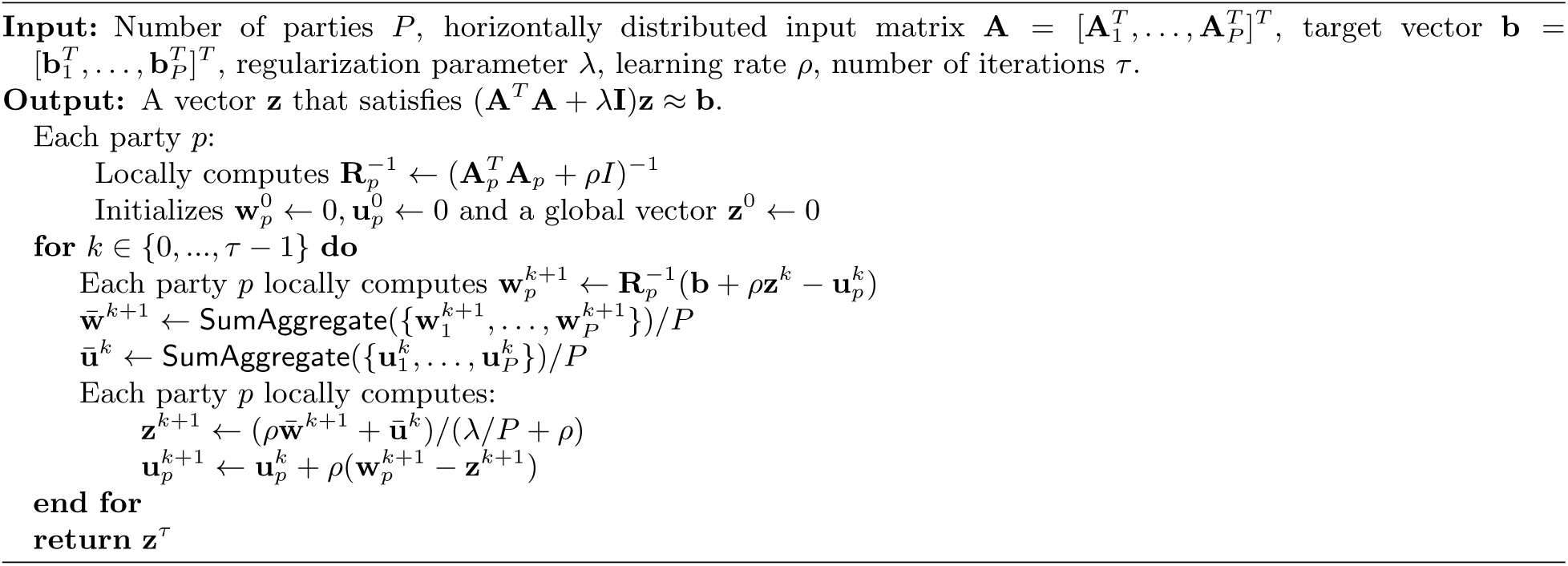

However, we note that our given 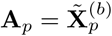 is not available in plaintext, as it is meant to be standardized and covariate-corrected based on the global matrix 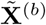. Therefore, although each party has access to their own raw genotype martix 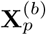, they are not able to precompute the following matrix shown in Algorithm 2 in plaintext:

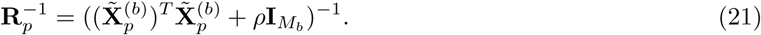

In SafeGENIE, we introduce a technique to resolve this issue by using the Woodbury matrix identity [39] to perform covariate correction in the computation of 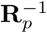 on the fly as follows. First, recall that

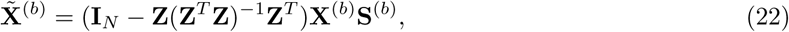

where **Z** is an N-by-C covariate matrix, and **S** represents a diagonal matrix with inverse standard deviations for each column of **X**^(*b*)^. We include an all-ones vector as a covariate in **Z**, which implicitly accounts for mean centering of **X**^(*b*)^. With one round of aggregation, we precompute a small *C*-by-*M*_*b*_ matrix

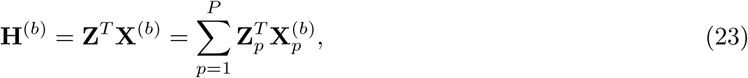

where each summand is computed locally using plaintext matrices then aggregated in an encrypted form. Next, noting that

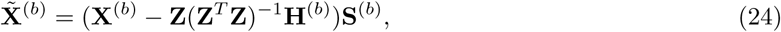

we are able to express 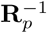 (Equation 21) as

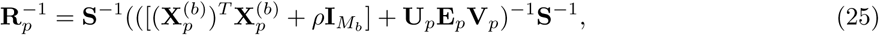

for some matrices 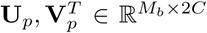 and **E**_*p*_ ∈ ℝ^2*C*×2*C*^ (see Appendix for full derivation; note the inner dimension of 2*C*). Finally, using the Woodbury identity and letting 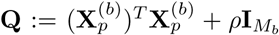 to simplify the notation, we can expand the inverse as

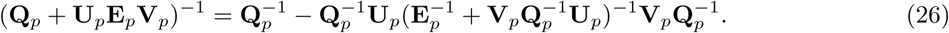

We have successfully reformulated the computation of 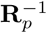 as one involving an *inverse of a plaintext matrix* **Q**_*p*_ and several matrix multiplications with a small inner dimension of 2*C*. Note that the new composite inverse matrix

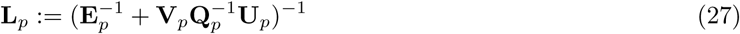

can be efficiently computed using secure MPC protocols, using the eigenfactorization routine introduced in prior work [13].

As a result, we can compute a key matrix 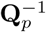 completely locally in plaintext, then use Equations 7 and 2 to compute the multiplication with 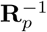 on the fly using the plaintext 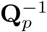. Note that this step is the only expensive matrix multiplication in the ADMM algorithm, and as such our reformulated ADMM offers significant reduction in computational cost. Moreover, we emphasize that, aside from the precomputation of **H**^(*b*)^ and 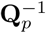, none of the matrix operations in our ADMM algorithm scales with the number of individuals in the dataset, and thus scales very efficiently to datasets with many samples, as our results show. Our final ADMM algorithm for ridge regression with covariates, leveraging the Woodbury identity technique, is presented in Algorithm 3 (changes with respect to the standard formulation are shown in blue).

### 3.6 Computational Complexity of SafeGENIE

SafeGENIE greatly reduces the runtime of a direct implementation of REGENIE in a distributed setting. By using our improved ADMM algorithm, we delegate large matrix inverse operations to be performed locally in plaintext. Since in practice the overhead of cryptographic operations greatly overshadows that of plaintext computation, in our complexity analysis we only consider homomorphic operations over the encrypted data.

The implementation of ADMM with our Woodbury optimization is separated into two components, the precomputation of a small matrix inverse in the Woodbury identity and the main ADMM iterations. For the precomputation of the **L**_*p*_ matrix (Equation 27), each party performs a *M*_*b*_-by-*M*_*b*_ matrix multiplication around 2*C* times where *C* is the number of covariates, where *M*_*b*_ is the blocksize. Each main iteration is dominated by the work of multiplying a *M*_*b*_-by-*M*_*b*_ plaintext matrix 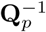twice with a ciphervector, combined with cipher-cipher multiplications with precomputed matrices **U**_*p*_, **L**_*p*_, and **V**_*p*_. Like in the plaintext setting, matrix vector multiplication in HE scales linearly with the size of the matrix. Since ridge regression in Level 0 is computed for each block (*B*), cross-validation fold (*K*), and regularization parameter (*R*), the complexity of Level 0 is *O*(*KBRT* ^2^ *τ*) where *τ* is the number of iterations in ADMM. Since *M* = *BM*_*b*_ by definition, this can be expressed as *O*(*M*_*b*_*KRM τ*). We note that this is a much better asymptotic runtime than naively using CGD for Level 0. Each iteration of CGD scales linearly with the size of the matrix being multiplied, which is *N*-by-*M*_*b*_ in our case. Therefore, the total runtime of Level 0 with CGD is *O*(*M*_*b*_*KBRN τ*), or equivalently *O*(*KRMN τ*), which is a factor of *N*/*M*_*b*_ larger than our ADMM method. For a dataset of *N* = 10^5^ and *M*_*b*_ = 10^3^, this amounts to a factor of 100 improvement using our ADMM solution, asymptotically.

#### Algorithm 3

Our Improved ADMM Algorithm for Ridge Regression with Covariates

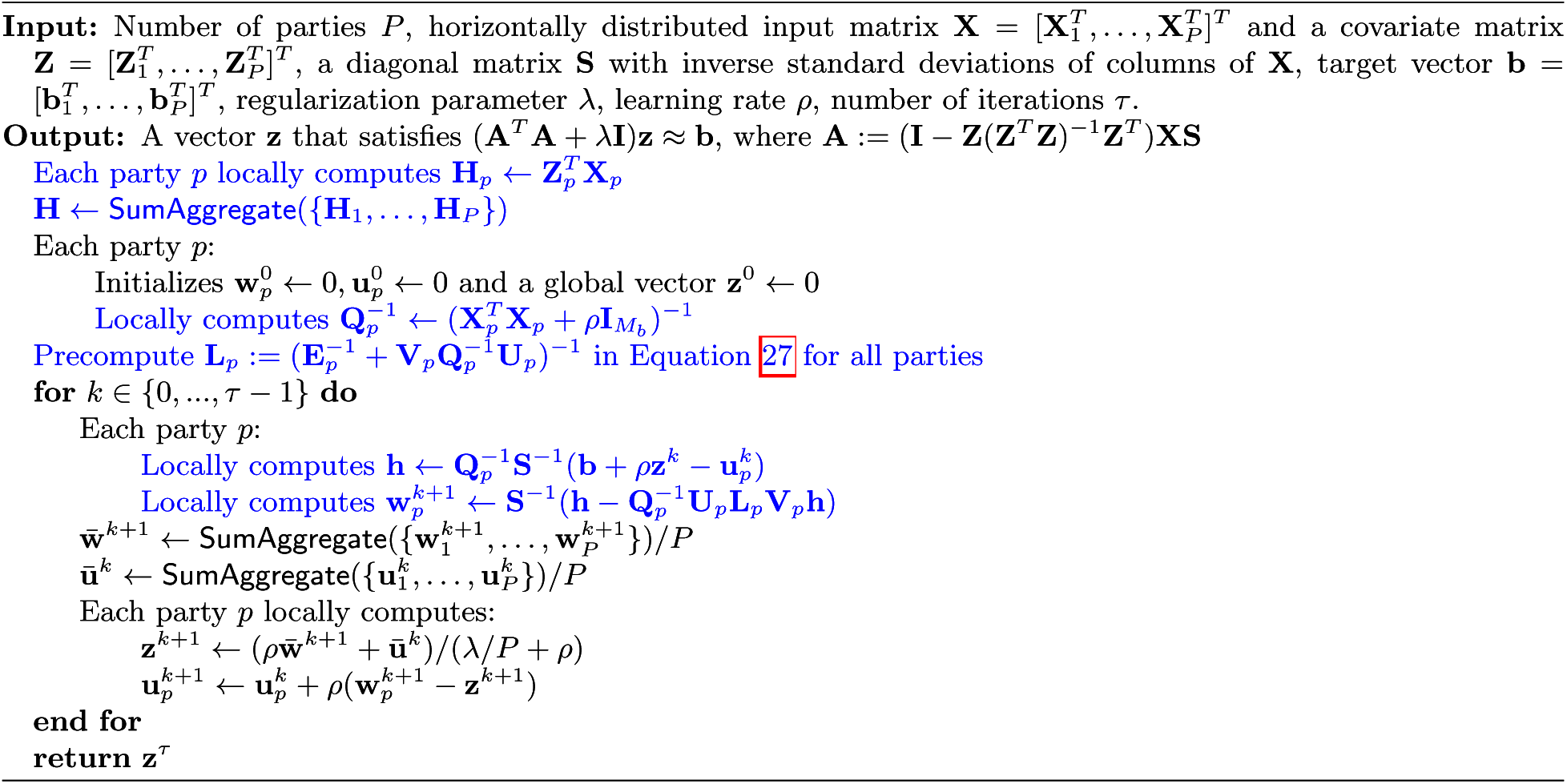

For Level 1, since we evaluate the CGD subroutine lazily without explicitly constructing the design matrix, we distribute the work in a way where each party multiplies their respective plaintext genotype matrix with a vector. Since the size of the matrix is *N*-by-*BR*, and there are *KR* different ridge regressions that must be performed, the runtime complexity of Level 1 is *O*(*KR*^2^*NB τ*), where *τ* is the number of iterations in CGD (typically 30), and *K* and *R* are small numbers (default values of 5 in REGENIE).

Lastly, to compute association statistics, much of the work can be done in plaintext. The chromosome-specific LOCO residual vectors (observed phenotypes minus the LOCO predictors of genetic effect) must be covariate corrected and then multiplied by the genotype matrix for the corresponding chromosome. This is equivalent to one matrix-vector multiplication between the full genotype matrix (plaintext) and a residual vector (ciphertext). While this leads to an asymptotic complexity of *O*(*NM*), in practice since only one such multiplication is needed and because cipher-plain multiplications are considerably more efficient than cipher-cipher multiplications, computing association statistics is the quickest of the three levels of computation.

### 3.7 Numerical Stability of SafeGENIE

There are several hyperparameters and statistics to take note of with SafeGENIE’s use of iterative methods for ridge regression that factor into the numerical stability and convergence time of SafeGENIE. Our ADMM algorithm for Level 0 depends on the hyperparameter *ρ*, which represents the step size used in the ADMM iterations. We have found that in practice, an effective *ρ* can be determined as a function of the number of individuals, since the genomic block size is typically chosen within a fixed range of 1K to 5K variants for REGENIE. In our experiments, we set *ρ* to the number of individuals in the local dataset divided by the number of cross-validation folds.

Another hyperparameter of interest for both CGD and ADMM is the number of iterations. The convergence of both methods are generally dependent on the condition number of the input matrix [37, 40] and thus rely heavily on the magnitude of the regularization parameter relative to the input matrix. Therefore, we implemented an adaptive approach that checks for convergence at regular intervals and terminates when a suitable solution has been obtained. We also employed a warm start strategy whereby solutions from a higher value of the regularization parameter is used to initialize the run with a lower value, which considerably improved convergence overall.

We also note that precision is challenging to maintain in general in secure computation protocols given their dependence on fixed-point representation of continuous numbers and the possibilities of numerical under/overflow. Throughout our algorithm, we carefully managed the range of data values by scaling intermediate results accordingly (e.g. dividing by 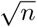 when computing the sum of *n* values for large *n*). As our results show, the combination of these strategies allow SafeGENIE to obtain accurate association analysis results.

### 3.8 Implementation Details

SafeGENIE is implemented using the distributed CKKS framework in Lattigo [33], which is an open source library in Golang for homomorphic encryption schemes. We extended the library by implementing secret sharing-based MPC functionalities based on our prior work [13]. In addition, SafeGENIE was implemented using Golang’s built in multi-threading framework and networking protocols, which allow interactive computation over encrypted data to be performed in a parallel manner across multiple machines. Our implementation also features efficient streaming pipelines for accessing blocks of the genotype matrix, which helps ensures that the memory usage of the program stays low regardless of the size of the dataset.

## 4 Experimental Results

### 4.1 SafeGENIE obtains accurate association statistics

To demonstrate the performance of SafeGENIE on a real GWAS dataset, we obtained a dataset of 9,178 East Asian lung cancer patients (5,054) and control individuals (4,053) from the dbGaP repository (accesion phs000716.v1.p1). After a quality control filter excluding individuals with missingness rate higher than 10%, we retained 9,098 individuals. We included in the analysis 378,482 single nucleotide polymorphisms (SNPs) that passed minor allele frequency (>0.1) and Hardy-Weinberg equilibirum (χ^2^ < 28.374) filters. We used a block size of 3000 SNPs for local ridge regressions in Level 0. For the association tests, we applied the standard leave-one-chromosome-out (LOCO) scheme, leaving out one chromosome at a time where the tested variant resides to correct for the background genetic effect.

For experimental setup, we created a set of three VM instances in the Google Cloud Platform, each with 128 RAM and 16 virtual CPUs, located in the same geographic zone. One VM served the role of a coordinating party for server-aided MPC routines, with the other two as main data holders participating in the collaborative LMM analysis. We split the GWAS data into two sets of individuals (4550 and 4548 individuals, respectively) and individually uploaded the corresponding genotype, phenotype, and covariate data to the two main parties’ VMs. We then executed the SafeGENIE program with point-to-point communication channels between pairs of parties for interactive steps of the protocol. For Level 0, we used additional sets of VMs to further parallelize the computation over the blocks, the results of which were later combined (and the reported runtime appropriately aggregated) before proceeding with the rest of the protocol. We used 12 threads on each machine to leverage parallelism in each step of the pipeline.

Figure 2 shows the resulting LMM-based association statistics from SafeGENIE compared against the output from running REGENIE (obtained from https://rgcgithub.github.io/regenie/) on a pooled dataset. The first two Manhattan plots show the genome-wide association signals are nearly identical between the two approaches, suggesting that SafeGENIE successfully replicates the analysis performed by REGENIE in a centralized setting. Note that SafeGENIE never has access to the whole data in one site; it only jointly analyzes distributed datasets in a secure manner using our cryptographic protocols. We observe that in this dataset a particularly strong association is identified for SNP rs2736100, which is associated with the *TERT* gene, a known cancer gene whose expression is associated with general increased cancer risk and is hypothesized to impair telomere maintenance [41]. Quantitatively measuring the agreement between the two outputs resulted in a correlation coefficient of 0.9845 (Figure 2C). We also observed a strong agreement in the genome-wide polygenic predictions of the phenotype (Figure 2D), which represent a key intermediate result in the LMM analysis.

**Figure 2:**
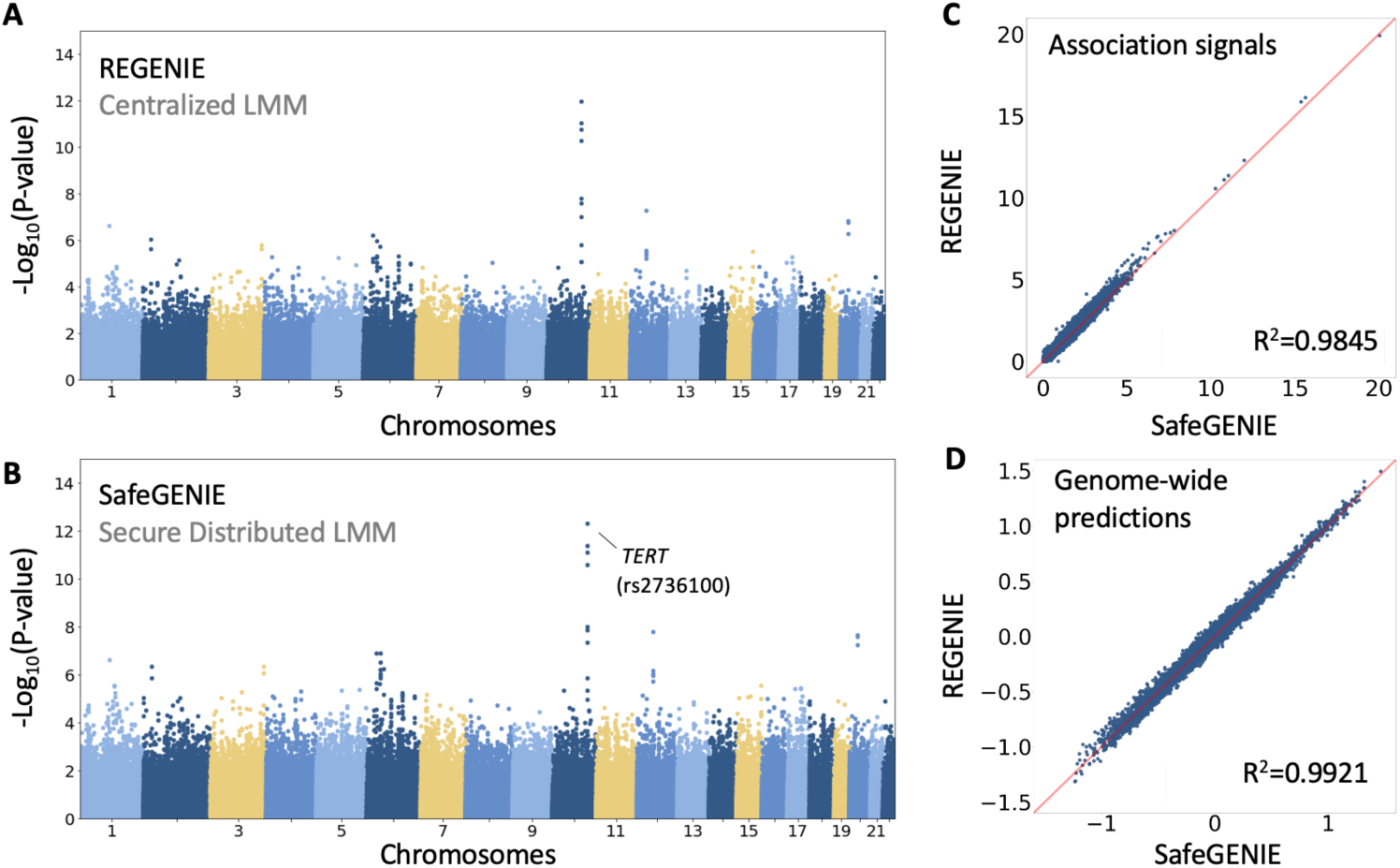
SafeGENIE closely reproduces REGENIE association statistics while securely analyzing distributed GWAS datasets. We evaluated SafeGENIE on a real lung cancer GWAS dataset including 9,098 individuals and 378,482 SNPs split between two parties. The Manhattan plot for SafeGENIE mirrors the Manhattan plot obtained from running the centralized REGENIE on the same lung cancer dataset (**A, B**). (**C**) Comparison of the negative log *p*-values for all variants in the dataset generated by REGENIE and by SafeGENIE. (**D**) Plot of the genome-wide phenotype prediction vectors obtained by both methods, which are provided as input to the association testing pipeline. Both show that the results from SafeGENIE are highly correlated to those of REGENIE with an *R*^2^ value over 0.98 in both plots.

**Figure 3:**
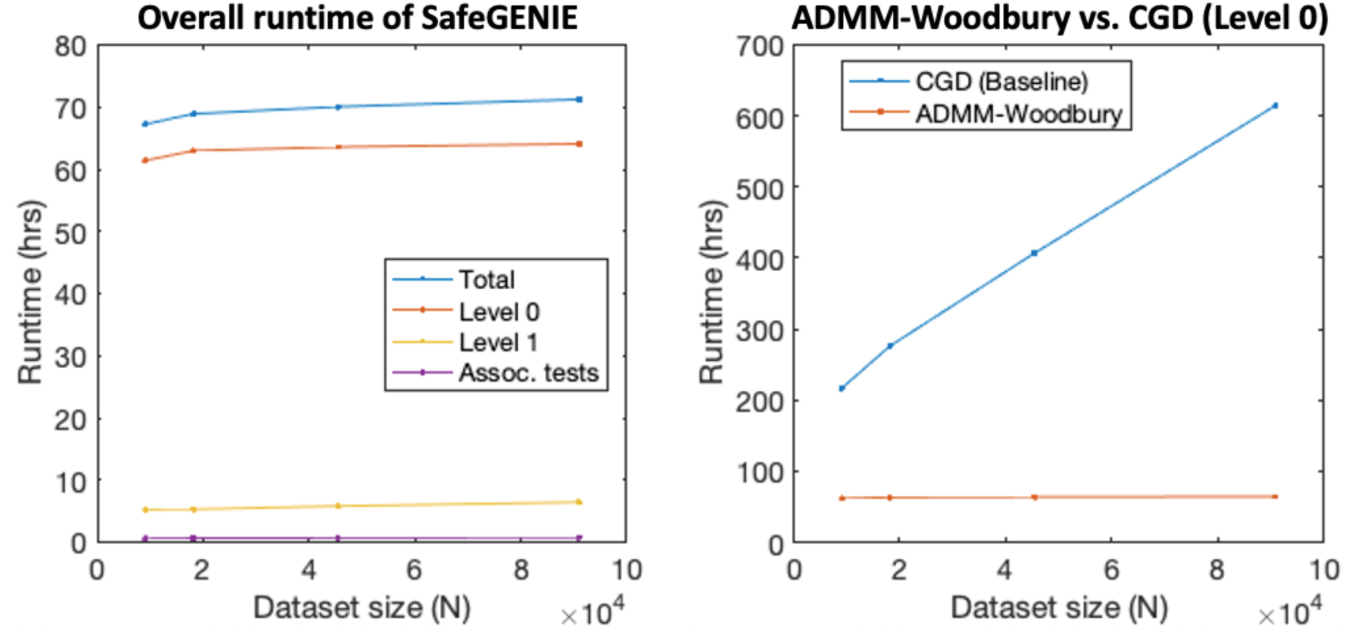
SafeGENIE achieves near-constant scalability with respect to runtime to larger datasets. We show the runtime of SafeGENIE on upsampled lung cancer datasets including up to 91K thousand individuals (ten times the original dataset). Runtimes shown are estimated based on a smaller number of blocks or iterations across which the computational load is expected to be identical. Measurements are based on a network of three co-located machines on Google Cloud with 12 cores each for parallelization. SafeGENIE’s runtime remains near-constant as the size of the dataset grows, enabled by the efficient distributed algorithms introduced in SafeGENIE, which maximize the use of local plaintext computation. Right subfigure shows the comparison of our ADMM-Woodbury algorithm for Level 0 with the baseline CGD algorithm (which we also newly developed for the secure distributed setting), demonstrating the improved asymptotic complexity of our approach. For the largest dataset, our approach achieves a 9.6-fold speedup over the baseline.

### 4.2 SafeGENIE efficiently scales to large datasets

The total runtime of SafeGENIE on the lung cancer dataset was 68.3 hours with a communication of 11 TB using 12 threads. Most of the runtime as well as communication can be attributed to Level 0 (61.4 hours and 11 TB; compared to 6.33 hours and 44.3 GB of Level 1), which fits many local ridge regression models across genomic blocks. We note that this step is embarrassingly parallel and thus can be sped up with more cores. In contrast, a baseline CGD solution for Level 0 without our ADMM-Woodbury algorithm, is estimated to take 216.75 hours of runtime (3.5 times slower) on 12 threads and 7.5 TB of communication (0.7 times less). Our runtime improvement is expected to be even greater at larger scales given our reduced dependence on *N* as described in Section 3.6.

We next evaluated the scalability of SafeGENIE on upsampled lung cancer datasets up to ten times its original size (91K individuals for the largest dataset) (Figure 2). We observed that the runtime of SafeGENIE is dominated by Level 0, whose runtime remained near-constant over the range of data sizes we tested. This incredible scalability is a result of the dimensionality reduction that is evident in our ADMM-Woodbury algorithm. Recall that to perform ridge regression over a genomic block, our algorithm calculates the inverse of a *M*_*b*_-by-*M*_*b*_ matrix in plaintext (where *M*_*b*_ is the block size) and perform all subsequent operations using this matrix, effectively removing dependence of runtime on the dataset size *N*. In contrast, the baseline conjugate gradient descent (CGD) solution for Level 0 ridge regressions (see Algorithm 1) grows linearly with *N*, resulting in a wider gap in performance compared to our algorithm for larger datasets (Figure 2); on the largest dataset, our ADMM-Woodbury solution achieves a factor of 9.6 speedup over CGD. Note that, while plaintext CGD is a widely used technique for ridge regression (also used in BOLT-LMM [21]), here we compared to our novel *secure distributed* implementation of CGD, which we used in Level 1. Since the majority of work in Level 0 consists of separate invocations of ridge regression on genomic blocks, this step can also be parallelized across multiple CPUs and machines to further reduce the runtime in practice. These results demonstrate the practical feasibility of SafeGENIE for large GWAS datasets including hundreds of thousands of individuals.

## 5 Discussion

We introduced SafeGENIE, a privacy-preserving and distributed approach to LMM association studies. Leveraging the insight that a recent stacked regression approach to LMM presents a path for efficient distributed computation, we developed efficient distributed algorithms for ridge regression with covariates for use as core routines in SafeGENIE. Our results show that SafeGENIE produces nearly identical association results compared to REGENIE, a centralized LMM algorithm, while demonstrating efficient runtime performance (in the order of days) with near-constant scalability to larger datasets. Directions for future work include evaluation on biobank-scale cohorts; increasing the robustness of SafeGENIE to a wide range of parameter settings (e.g. including extreme values of variance estimates); and further developing methods to support other types of association tests based on LMM beyond the quantitative trait model addressed in this work. We expect the insights introduced in our work on how to design efficient distributed algorithms for secure computation to be of broad interest to enhancing privacy in other genomic analysis workflows.

## Footnotes

1 This manuscript has been accepted for presentation in the Genome Privacy and Security Workshop at RECOMB 2022.

