## Supplementary Materials for "Secure and federated linear mixed model association tests"

### Appendix

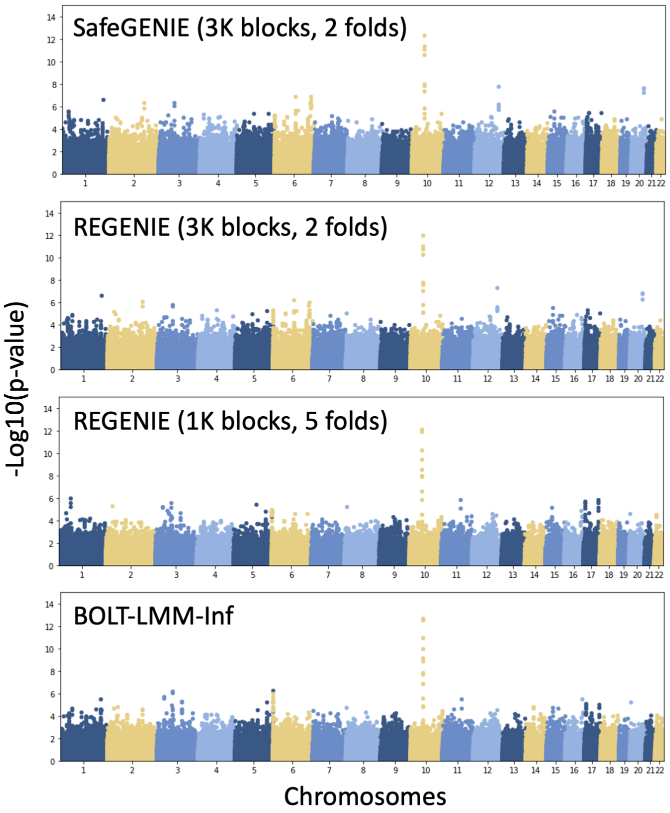


Supplementary Figure 1: **Comparable performance of REGENIE and BOLT-LMM-Inf on the lung cancer dataset.** We compared the association results of REGENIE on the lung cancer dataset with BOLT-LMM, a popular method for LMM analysis. Top two figures reproduce our results in Figure 2, which are based on a parameter setting with 3K variants per block and 2 folds. These results demonstrate that SafeGENIE closely matches the performance of REGENIE. Bottom two figures show similar results between BOLT-LMM-Inf (based on the infinitesimal model analogous to REGENIE) and REGENIE with 1K variants per block and 5 folds for cross-validation. Because our dataset includes a moderate number of individuals (9098) and SNPs (378K), we expect smaller block sizes and more folds to lead to better approximation performance in general for REGENIE, which explains the increased discrepancy between BOLT-LMM and REGENIE in our parameter setting. Note that SafeGENIE can run with any suitable choice of parameters for a given dataset to emulate the centralized REGENIE analysis in a distributed setting.

#### Derivation of Improved ADMM based on the Woodbury Identity

To better exploit the power of ADMM, we aim to expand the following inverse which denotes the inverse matrix that is computed locally by each party.

$$\mathbf{R}_p^{-1} = ((\tilde{\mathbf{X}}_p^{(b)})^T \tilde{\mathbf{X}}_p^{(b)} + \rho \mathbf{I}_{M_b})^{-1}. \quad (1)$$

The Woodbury Matrix identity allows for the expansion of inverses into a new inverse with a small rank one update if the initial matrix is of the following form

$$(\mathbf{Q} + \mathbf{UEV})^{-1} = \mathbf{Q}^{-1} - \mathbf{Q}^{-1}\mathbf{U}(\mathbf{E}^{-1} + \mathbf{VQ}^{-1}\mathbf{U})^{-1}\mathbf{VQ}^{-1}. \quad (2)$$

We can begin by expanding $\tilde{\mathbf{X}}_p^{(b)}$ into its components. Namely, we know that

$$\tilde{\mathbf{X}}^{(b)} = (\mathbf{X}^{(b)} - \mathbf{Z}(\mathbf{Z}^T\mathbf{Z})^{-1}\mathbf{H}^{(b)})\mathbf{S}^{(b)}. \quad (3)$$

Therefore,

$$\mathbf{R}_p^{-1} = (((\mathbf{X}^{(b)} - \mathbf{Z}(\mathbf{Z}^T\mathbf{Z})^{-1}\mathbf{H}^{(b)})\mathbf{S}^{(b)})^T(\mathbf{X}^{(b)} - \mathbf{Z}(\mathbf{Z}^T\mathbf{Z})^{-1}\mathbf{H}^{(b)})\mathbf{S}^{(b)} + \rho\mathbf{I}_{M_b})^{-1}, \quad (4)$$

After flipping the inner transpose, we get the following

$$\mathbf{R}_p^{-1} = (\mathbf{S}^{(b)}((\mathbf{X}^{(b)})^T - (\mathbf{H}^{(b)})^T(\mathbf{Z}^T\mathbf{Z})^{-1}\mathbf{Z}^T)(\mathbf{X}^{(b)} - \mathbf{Z}(\mathbf{Z}^T\mathbf{Z})^{-1}\mathbf{H}^{(b)})\mathbf{S}^{(b)} + \rho\mathbf{I}_{M_b})^{-1}, \quad (5)$$

After distribution the expression becomes

$$\mathbf{R}_p^{-1} = ((\mathbf{X}^{(b)})^T\mathbf{X}^{(b)} + \rho\mathbf{I}_{M_b} - (\mathbf{H}^{(b)})^T(\mathbf{Z}^T\mathbf{Z})^{-1}\mathbf{Z}^T\mathbf{X}^{(b)} - (\mathbf{X}^{(b)})^T(\mathbf{Z}^T\mathbf{Z})^{-1}\mathbf{H}^{(b)} + (\mathbf{H}^{(b)})^T(\mathbf{Z}^T\mathbf{Z})^{-1}\mathbf{Z}^T\mathbf{Z}(\mathbf{Z}^T\mathbf{Z})^{-1}\mathbf{H}^{(b)})^{-1},$$

We then consolidate two terms together.

$$\mathbf{R}_p^{-1} = ((\mathbf{X}^{(b)})^T\mathbf{X}^{(b)} + \rho\mathbf{I}_{M_b} - (\mathbf{X}^{(b)})^T(\mathbf{Z}^T\mathbf{Z})^{-1}\mathbf{H}^{(b)} - (\mathbf{H}^{(b)})^T(\mathbf{Z}^T\mathbf{Z})^{-1}(\mathbf{Z}^T\mathbf{X}^{(b)} - \mathbf{Z}^T\mathbf{Z}(\mathbf{Z}^T\mathbf{Z})^{-1}\mathbf{H}^{(b)}))^{-1},$$

For simplicity of notation let $\mathbf{J} = \mathbf{Z}^T\mathbf{Z}(\mathbf{Z}^T\mathbf{Z})^{-1}\mathbf{H}^{(b)}$. We can then simply the expression to

$$\mathbf{R}_p^{-1} = ((\mathbf{X}^{(b)})^T\mathbf{X}^{(b)} + \rho\mathbf{I}_{M_b} - (\mathbf{X}^{(b)})^T(\mathbf{Z}^T\mathbf{Z})^{-1}\mathbf{H}^{(b)} - (\mathbf{H}^{(b)})^T(\mathbf{Z}^T\mathbf{Z})^{-1}(\mathbf{Z}^T\mathbf{X}^{(b)} - \mathbf{J}))^{-1},$$

Let $M_b$ denote blocksize and C denote the number of covariates. Now let

$$\mathbf{U} = \begin{bmatrix}\mathbf{H}^{(b)} & (\mathbf{X}^{(b)})^T\mathbf{Z}\end{bmatrix},$$

which is a $M_b \times 2C$ matrix

$$\mathbf{E} = \begin{bmatrix}\mathbf{Z}^T\mathbf{Z} & 0 \\ 0 & \mathbf{Z}^T\mathbf{Z}\end{bmatrix}^{-1},$$

which is a $2C \times 2C$ matrix

$$\mathbf{V} = \begin{bmatrix}\mathbf{Z}^T\mathbf{X}^{(b)} - \mathbf{J} \\ \mathbf{H}^{(b)}\end{bmatrix},$$

which is a $2C \times M_b$ matrix

$$\mathbf{R}_p^{-1} = \mathbf{S}^{-1}(([\mathbf{X}_p^{(b)})^T\mathbf{X}_p^{(b)} + \rho\mathbf{I}_{M_b}] + \mathbf{UEV})^{-1}\mathbf{S}^{-1}, \quad (6)$$

Now, using the Woodbury Matrix Identity, we can set $\mathbf{Q} = (\mathbf{X}_p^{(b)})^T\mathbf{X}_p^{(b)} + \rho\mathbf{I}_{M_b}$ and expand the inverse

$$\mathbf{R}_p^{-1} = \mathbf{S}^{-1}[\mathbf{Q}^{-1} - \mathbf{Q}^{-1}\mathbf{U}(\mathbf{E}^{-1} + \mathbf{VQ}^{-1}\mathbf{U})^{-1}\mathbf{VQ}^{-1}]\mathbf{S}^{-1}, \quad (7)$$

Note that $(\mathbf{E}^{-1} + \mathbf{VQ}^{-1}\mathbf{U})^{-1}$ is a very small inverse matrix which is $2C \times 2C$ in size. This allows us to compute this inverse explicitly without much issue. The above expression is mathematically equivalent to $\mathbf{R}_p^{-1}$ and thus can be substituted for any instance of $\mathbf{R}_p^{-1}$ in the original ADMM algorithm. This substitution is what becomes Algorithm 3—our improved ADMM algorithm for ridge regression with covariates.

#### Association testing

Here we outline how association testing is performed in SafeGENIE. For each variant $g$, we wish to calculate

$$s := \frac{\tilde{g}^T \hat{y}^*_{\text{resid,LOCO}}}{\hat{\sigma}_e \cdot \sqrt{\tilde{g}^T \tilde{g}}}.$$

Note that

$$\tilde{g} = \sigma_g^{-1} P g,$$

where $P = (I - Z(Z^T Z)^{-1} Z^T)$. Since we include a column of ones in the covariate matrix $Z$ for mean shift correction, we do not need to apply mean correction in advance to $g$. Also, since the scaling factor $\sigma_g^{-1}$ cancels between the denominator and the numerator, we can set it to one without affecting the result, i.e. let $\tilde{g} = Pg$. Thus, we can express the target quantity as

$$s = \frac{g^T P \hat{y}^*_{\text{resid,LOCO}}}{\hat{\sigma}_e \cdot \sqrt{g^T P g}}$$

We first directly calculate $\hat{y}^*_{\text{resid,LOCO}}$ and $\hat{\sigma}_e$. Then, we compute $g^T P \hat{y}^*_{\text{resid,LOCO}}$ by first computing

$$w := P \hat{y}^*_{\text{resid,LOCO}}$$

then computing

$$\tilde{g}^T \hat{y}^*_{\text{resid,LOCO}} = g^T w.$$

Note that $w$ is computed once per chromosome and shared across all variants.

Next we consider $g^T P g$. We have access to a matrix $R$ such that $RR^T = (Z^T Z)^{-1}$, using which we can express $P = I - ZRR^T Z^T$. Define

$$u := R^T Z^T g.$$

Then we can compute

$$g^T P g = g^T g - u^T u.$$

To further consolidate the computation above, we can take the following approach. We first compute

$$w := P \hat{y}^*_{\text{resid,LOCO}}.$$

Then, we take a single pass over the input genotype matrix to compute the following terms for each column $g$:

$$g^T g,\ Z^T g,\ w^T g.$$

Note that the first two are computed in plaintext and easily converted to secret shares, where each party sets their own share to the computed plaintext value such that the implicit sum of shares simply becomes the overall sum. The third term involves a ciphervector $w$ and thus can be treated as ciphervector-plainmatrix multiplication.

Given these terms, we can compute $u$ using the secret shares, then compute $g^T P g$ as well as its inverse square-root using the secret shares as well. Afterwards, we need only multiply the numerator with the inverse of the denominator (taking $\hat{\sigma}_e$ into account) and obtain the final statistics.
